# Differential effects of corticosteroids and anti-TNF on tumor-specific immune responses - implications for the management of irAEs

**DOI:** 10.1101/437830

**Authors:** Arianna Draghi, Troels Holz Borch, Haja Dominike Radic, Christopher Aled Chamberlain, Aishwarya Gokuldass, Inge Marie Svane, Marco Donia

**Affiliations:** Center for Cancer Immune Therapy, Department of Hematology, Herlev and Gentofte Hospital, University of Copenhagen, Herlev, Denmark; Department of Oncology, Herlev and Gentofte Hospital, University of Copenhagen, Herlev, Denmark

**Keywords:** Immune-related adverse events, Corticosteroids, Anti-TNF, *In vitro* tumor-killing, Cancer immunotherapy

## Abstract

Up to 60% of patients treated with cancer immunotherapy develop severe or life threatening immune-related adverse events (irAEs). Immunosuppression with high doses of corticosteroids or, in refractory cases, with tumor necrosis factor (TNF) antagonists, are the mainstay of treatment for irAEs. It is currently unknown what is the impact of corticosteroids and anti-TNF on the activity of antitumor T cells. In this study, the influences of clinically relevant doses of dexamethasone (corresponding to an oral dose of 10 to 125 mg prednisolone) and infliximab (anti-TNF) on the activation and killing ability of tumor-infiltrating lymphocytes (TILs) was tested *in vitro*. Overall, dexamethasone at low or intermediate/high dose impaired the activation (respectively −46% and −62%) and tumor-killing ability (respectively −48% and −53%) of tumor-specific TILs. In contrast, a standard clinical dose of infliximab only had a minor effect on T cell activation (−20%) and tumor killing (−10%). A brief resting following exposure to dexamethasone was sufficient to rescue the *in vitro* activity of TILs. In conclusion, clinically-relevant doses of infliximab only influenced to a lesser extent the activity of tumor-specific TILs *in vitro*, whereas even low doses of corticosteroids markedly impaired the antitumor activity of TILs. These data support steroid-sparing strategies and early initiation of anti-TNF for the treatment of irAEs in immuno-oncology.

## Manuscript

### Introduction

Recent advances in cancer immunotherapy have yielded impressive long-lasting response in patients affected by metastatic melanoma and other cancers ^1^. Nevertheless, up to 90% ^2^ of patients treated with immune checkpoint inhibitors (ICI) can develop immune-related adverse events (irAEs) affecting most frequently skin, colon, endocrine glands, liver and lungs ^3, 4^. The physiological role of immune checkpoints is to maintain self-tolerance and control inflammation ^5^, thus it is not surprising that any interference in immune checkpoint signaling may lead to immune-mediated tissue damage. IrAEs can manifest at any time point during immunotherapeutic treatment and their late recognition or inappropriate management may result in life threatening consequences ^4, 6^. Prompt diagnosis and adequate treatment are highly warranted.

The first line clinical management for most irAEs ≥ grade 3 or intolerable/persisting ≥ grade 2 according to the Common Terminology Criteria for Adverse Events (CTCAE) scale, is based on the use of systemic corticosteroids ^3, 6^. International consensus guidelines recommend patients with steroid-refractory irAEs ≥ grade 3 to be promptly treated with second-line immunosuppressive agents, including anti-tumor necrosis factor (TNF) agents such as infliximab ^3, 7, 8^, and having their ICI to be discontinued. Although the number of patients treated with ICI is increasing dramatically, the current level of evidence on which these guidelines are based is low and there are no randomized trials comparing different strategies. Recently, other drugs such as vedolizumab were introduced in the second-line treatment of irAEs, but their role is more controversial ^9, 10^. Previous studies investigating the outcome of patients treated with steroids for the management of anti-CTLA-4 induced irAEs did not show a worse survival outcome of the steroid-treated group in melanoma ^11^, however baseline steroids diminished the survival benefits of PD-1/PD-L1 antibodies in lung cancer ^12^. Thus, it is still unclear whether systemic immunosuppression may jeopardize the efficacy of the adaptive immune system in fighting cancer or whether distinct drugs can mediate differential effects on auto-immune and anti-cancer responses. In this study, we aimed to address this issue by evaluating how clinically relevant doses of dexamethasone (a standard corticosteroid) or infliximab can modulate the activation and killing ability of tumor-infiltrating lymphocytes (TILs) *in vitro*.

## Materials and Methods

### Immune cells and tumors

TILs were isolated and expanded from metastatic melanoma lesions. All the samples were obtained from patients undergoing tumor resection or baseline core-biopsies for enrollment in clinical trials. Written informed consent was provided by all patients. All procedures were conducted in accordance with the Declaration of Helsinki and Good Clinical Practice. TILs were established *in vitro* with a two-step process; the initial expansion to produce “young TILs” and the rapid expansion (REP) to produce REP TILs, as previously described in detail ^13^. REP TILs were used for all the analyses on TILs. Autologous tumor cell lines were established as described elsewhere ^14^ using the same tumor lesion from which the TILs were generated. All the cell lines were generated internally and not otherwise authenticated. Mycoplasma testing was not performed and the number of passages between collection and use in the described experiments was <10. Tumor-TIL pairs with known high reactivity (at least 10% of reactive CD8+ T cells) were selected for these analyses. Peripheral blood mononuclear cells (PBMCs) were isolated with gradient-centrifugation from blood samples collected from patients treated in the trial NCT00937625 at least 2 years following immunotherapy. Tumor-PBMCs pairs with known reactivity (>1% tumor-reactive T cells in the blood) were selected for these analyses.

### Assays of T cell activation and T cell-mediated tumor killing

Tumor-specific immune activation and tumor-killing were assessed with co-culture assays of TILs or PBMCs and autologous tumor cells in the presence of different concentrations of water soluble dexamethasone (D2915, Sigma-Aldrich/Merck KGaA, Darmstadt, Germany), infliximab (EU/1/13/854/001, Hospira, Hurley, UK) and mouse IgG1 kappa isotype control (16-4714-82, Thermo Fisher Scientific, Waltham, MA, USA). Increasing doses of steroids, including 0.01 μM, 0.02 μM, 0.04 μM and 0.1 μM (respectively equivalent to the maximum free prednisolone plasma level reached following repeated administration of approximately 10 mg, 25 mg, 50 mg and 125 mg oral prednisolone ^15, 16^) were chosen, whereas infliximab was used at a fixed dose (10 μg/ml), corresponding to a level slightly above the highest range of steady-state concentrations for the standard infusion dose of 5 mg/kg ^17^. Experiments in the presence of dexamethasone, infliximab or isotype control were conducted by adding the drugs directly in the culture wells at the beginning of the assay. Withdrawal experiments were conducted by culturing TILs in standard complete TIL media containing IL-2 (as described in ^14^) for 4 days with or without dexamethasone and followed by the T cell assay without dexamethasone (short withdrawal); additional experiments were conducted after removing dexamethasone from the culture media and additional 3 days of resting. T cell activation was evaluated with flow cytometry at different time points, and it was defined as the percentage of T cells staining positive for the activation marker CD137 (4-1BB) minus control. The antibodies used for flow cytometry were: CD3 FITC (345764, BD Biosciences, San Jose, CA, USA), CD56 PE (345812, BD Biosciences, San Jose, CA, USA), CD8 PerCP (345774, BD Biosciences, San Jose, CA, USA), CD4 BV510 (562970, BD Biosciences, San Jose, CA, USA), CD137 APC (309810, BioLegend, San Diego, CA, USA) and Live/Dead Fixable Dead Cell Stain Near-IR (L34976, Thermo Fisher Scientific, Waltham, MA, USA). The tumor-specific killing ability of TILs was evaluated in the presence of different concentration of dexamethasone, infliximab or IgG1 isotype in the xCELLigence system RTCA DP real-time cell analyzer (Bundle 00380601050, Analyzer 05469759001, Control Unit 05454417001, ACEA Biosciences Inc. San Diego, CA) on E-plate 16 plates (05469830001, ACEA Biosciences Inc. San Diego, CA, USA), according to the manufacturer’s instructions.

### Statistical Analyses

Statistical tests were conducted using a paired Wilcoxon signed rank test. Graphs and statistical tests were generated using Graphpad Prism 5. Negative killing values were converted to 0.01% for statistical analyses and generation of figures. All values were expressed as median. Tumor-specific T cell activation in presence of IgG1 isotype (10 μg/ml) differed less than ±10% from tumor-specific T cell activation in absence of any reagent. Therefore, tumor-specific T cell activation in presence of both dexamethasone and infliximab was compared to the activation in the absence of any reagent.

## Results and Discussion

Overall, tumor-specific activation of CD8^+^ TILs after 8 hours of stimulation with autologous tumor cells was reduced by 46% and 62% with dexamethasone 0.01 μM and 0.1 μM, respectively (n=8; both p<0.01). In contrast, infliximab reduced CD8^+^ T cells activation by only 20% (n=8; p<0.01 versus control, p<0.05 versus dexamethasone 0.01 μM, p<0.01 versus dexamethasone 0.1 μM) (Figure 1A and 1B). This impaired T cell activation was maintained over time (Figure 1C). Similar results were obtained comparing the activation of CD8^+^ peripheral blood lymphocytes (PBLs) following recognition of autologous tumors from two patients (Figure 1D). Importantly, even high doses of dexamethasone did not abrogate T cell activation (Figure 1A, 1B and 1C). The tumor-killing ability of TILs at 24 hours was impaired by 48% and 53% respectively by dexamethasone 0.01 μM and dexamethasone 0.04 μM (n=6; both p<0.05) (Figure 2A). In contrast, infliximab reduced tumor killing by only 10% at 24 hours (n=6; p=n.s. versus control, p<0.05 versus dexamethasone 0.01 μM or dexamethasone 0.04 μM) (Figure 2A). As for T cell activation, this impaired T cell-mediated tumor killing was maintained over time (Figure 2B and 2C) and very high doses of steroids impaired but did not abrogate completely tumor killing (Figure 2A, 2B and 2C). Similar doses of dexamethasone, as used in previous experiments, did not have a detectable impact on the growth of tumor cells (Figure 2D). Only an extremely high, and not clinically relevant, dose of dexamethasone impaired tumor cell growth (Figure 2D). Following exposure to a dose of dexamethasone (0.02 μM for 4 days) which was expected to impair the activation and killing of >50% based on results presented in Figure 1 and Figure 2, 3 days resting after withdrawal of dexamethasone was sufficient to rescue the activation (Figure 3A) and killing ability (Figure 3B and 3C) of TILs, while a short withdrawal (only during the assay) did not appear to rescue fully T cell activation (Figure 3A) and killing (Figure 3C).

**Figure 1.**
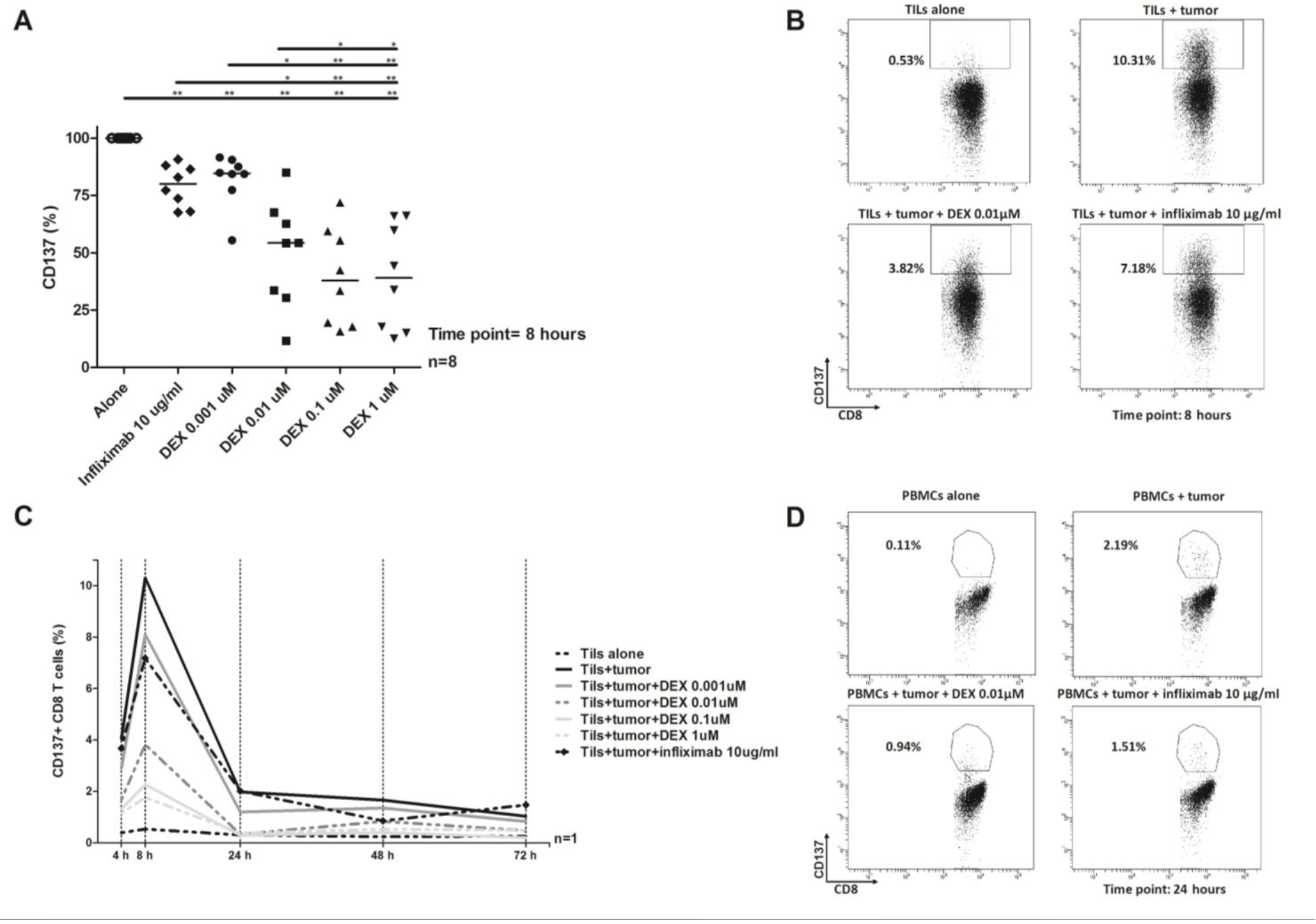
Modulation of CD8^+^ tumor-specific T cells activation by dexamethasone (DEX) or infliximab. (A) The panel shows the activation of CD8^+^ tumor-infiltrating lymphocyte (TILs) following 8 hours of co-culture with autologous tumor cells (n = 8). Tumor-specific activation in presence of dexamethasone 0.01 μM, dexamethasone 0.1 μM and 10 μg/ml infliximab was reduced by median 46%, 62%, and 20%, respectively. (B) FACS plots showing antitumor reactivity (CD137 positive CD8^+^ T cells) in TILs from a single exemplary donor following 8 hours of co-culture with autologous tumor cells. (C) The panel shows the activation of CD8^+^ TILs from a single exemplary donor at different time points after co-culture with autologous tumor cells. (D) FACS plots showing anti-tumor reactivity (CD137 positive CD8^+^ T cells) in CD8^+^ peripheral blood lymphocytes from a single exemplary donor following 24 hours of co-culture with autologous tumor cells. *p<0.05; **p<0.01

**Figure 2.**
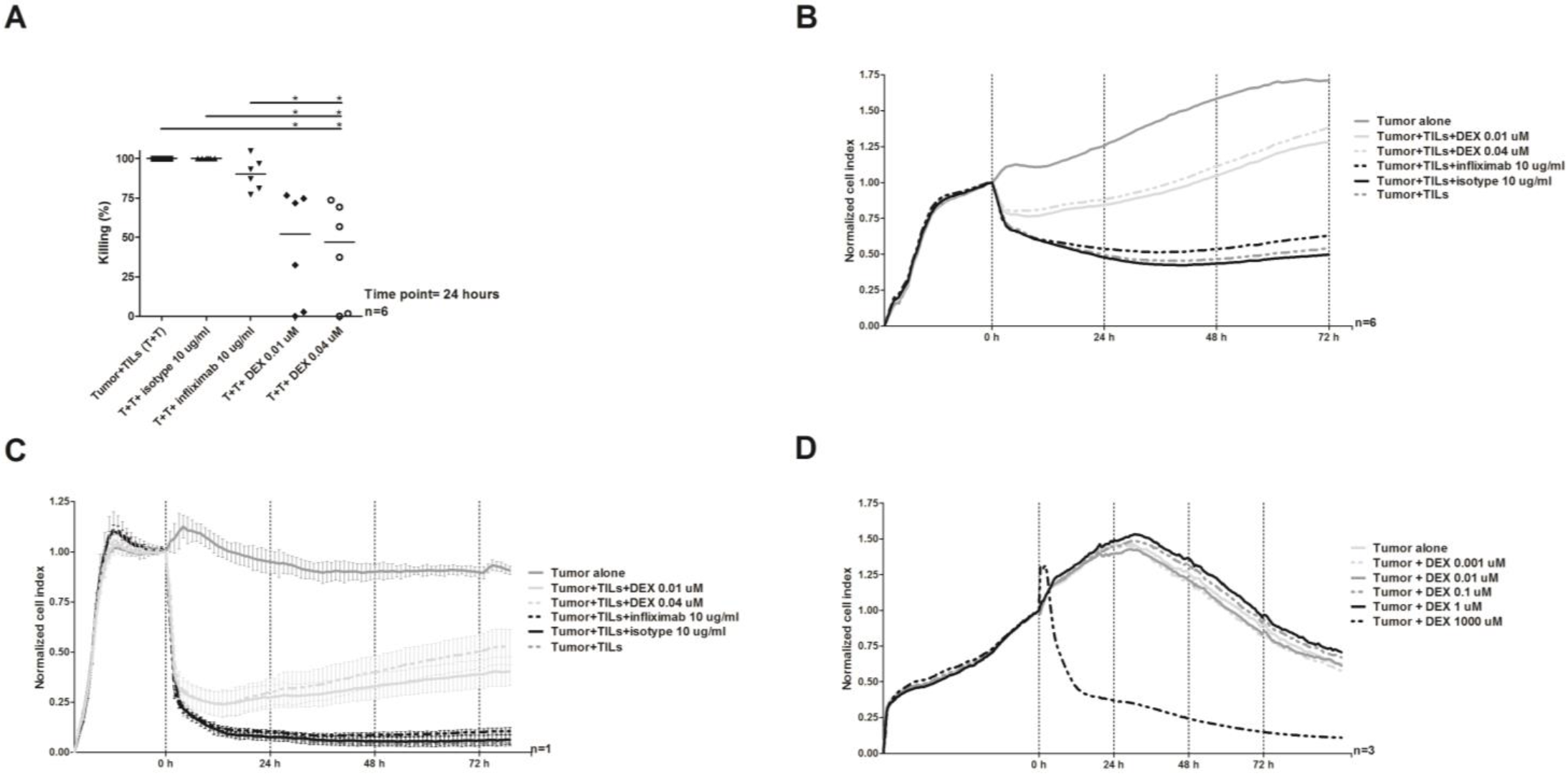
Modulation of tumor-infiltrating lymphocyte (TIL) killing by dexamethasone (DEX) or infliximab. (A) The panel shows the killing ability of TILs following 24 hours of co-culture with autologous tumor cells (n = 6). Tumor-specific killing in presence of dexamethasone 0.01 μM, dexamethasone 0.04 μM and 10 μg/ml infliximab was reduced by median 48%, 53% and 10%, respectively. (B) The panel shows the killing ability of TILs at different time points during co-culture with autologous tumor cells (n = 6). (C) The panel shows the killing ability of TILs from a single exemplary donor at different time points during co-culture with autologous tumor cells. (D) The panel shows the growth of tumor cells in presence of dexamethasone at different time points (n = 3). In (B) and (D), curves show the mean value of distinct samples. *p<0.05

**Figure 3.**
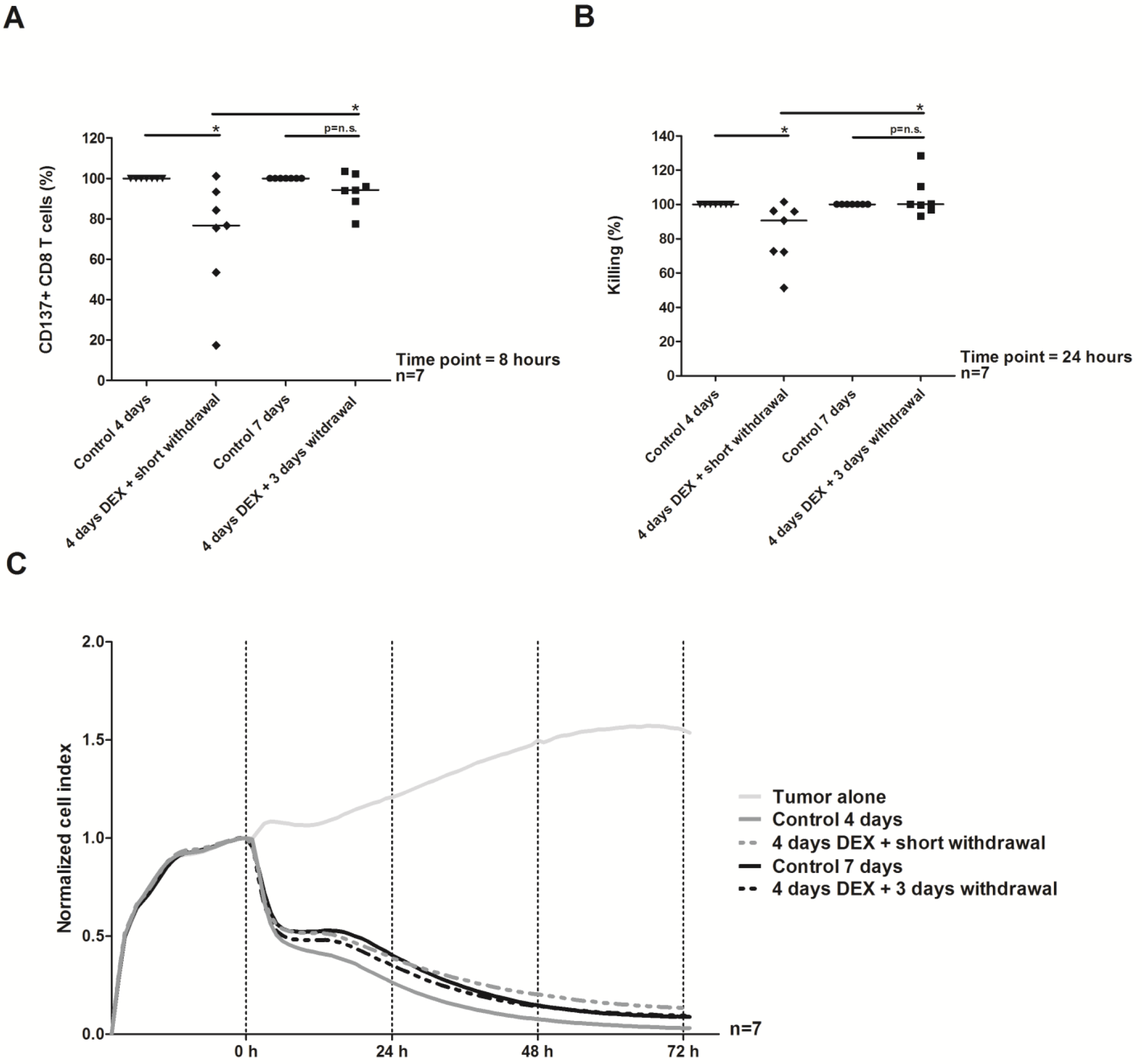
Modulation of tumor-infiltrating lymphocyte (TIL) response after dexamethasone (DEX) withdrawal. (A) The panel shows the activation of CD8^+^ TILs following 8 hours of co-culture with autologous tumor cells (n = 7). (B) The panel shows the killing ability of TILs following 24 hours of co-culture with autologous tumor cells (n=7). (C) The panel shows the killing ability of TILs at different time points during co-culture with autologous tumor cells (n = 7). In (C), curves show the mean value of distinct samples. *p<0.05

Prospective clinical studies on how immunosuppressive and immunomodulating drugs may influence the clinical efficacy of cancer immunotherapy are currently lacking. The use of steroids in patients with cancer is much broader than the management of irAEs, as steroids are also the mainstay of symptomatic treatment for brain metastases. Here we show that low to moderate doses of steroids had a negative impact on the activation and killing ability of TILs *in vitro*, but even high doses of steroids did not completely abrogate both activation and killing; importantly, steroid withdrawal may restore the activity of TILs. Noteworthy, anti-TNF antibodies only had a minor effect on the activity of TILs. In conclusion, these data support early use of anti-TNF for the treatment of serious irAEs and warrant prospective clinical studies to compare first line strategies for the management of irAEs. Of particular interest, a recent preclinical study by Bertrand et al.^18^, shows that treatment with ICI can be potentiated by TNF blockade. Prospective clinical testing of baseline steroids on the efficacy of ICI in patients with melanoma metastatic to the brain is already ongoing (NCT03563729).

## Acknowledgements

Marta Español-Rego (Department of Immunology-CDB, Hospital Clinic, IDIBAPS, Barcelona, Spain) is acknowledged for assistance in performing selected experiments. These works were supported by a research grant to Inge Marie Svane from The Danish Cancer Society, Knæk Cancer campaign.

## Availability of data and material

Additional data obtained during the current study or source materials are available from the corresponding author upon reasonable request.

## Abbreviations

CTCAE: Common Terminology Criteria for Adverse Events
ICI: Immune checkpoint inhibitors
irAE: Immune-related adverse event
PBL: Peripheral blood lymphocyte
PBMC: Peripheral blood mononuclear cell
REP: Rapid expansion
TIL: Tumor-infiltrating lymphocyte
TNF: Tumor necrosis factor

